# PGR5 is required to avoid photosynthetic oscillations during light transitions

**DOI:** 10.1101/2023.09.21.558795

**Authors:** Gustaf E. Degen, Federica Pastorelli, Matthew P. Johnson

## Abstract

The production of ATP and NADPH by the light reactions of photosynthesis and their consumption by the Calvin-Benson-Bassham (CBB) cycle and other downstream metabolic reactions requires careful regulation. Environmental shifts perturb this careful balance, leading to photo-oxidative stress and losses in CO_2_ assimilation. Imbalances in the production and consumption of ATP and NADPH manifest themselves as transient instability in the chlorophyll fluorescence, P700, electrochromic shift and CO_2_ uptake signals recorded on leaves. These oscillations can be induced in wild-type plants by sudden shifts in CO_2_ concentration or light intensity, however mutants exhibiting increased oscillatory behaviour have yet to be reported. This has precluded an understanding of the regulatory mechanisms employed by plants to suppress oscillations. Here we show that the Arabidopsis *pgr5* mutant, which is deficient in PGR5-dependent cyclic electron transfer (CET), exhibits increased oscillatory behaviour. In contrast, mutants lacking the NDH-dependent CET are largely unaffected. The absence of oscillations in the *hope2* mutant, which like *pgr5*, lacks photosynthetic control and exhibits high ATP synthase conductivity, ruled out loss of these photoprotective mechanisms as causes. Instead, we observed slower formation of proton motive force and by inference ATP synthesis in *pgr5* following environmental perturbation, leading to the transient reduction of the electron transfer chain and photosynthetic oscillations. PGR5-dependent CET therefore plays a major role in damping the effect of environmental perturbations on photosynthesis to avoid losses in CO_2_ fixation.

## Introduction

The light reactions of photosynthesis supply ATP and NADPH to the Calvin-Benson-Bassham (CBB) cycle, the site of CO_2_ fixation. The CBB cycle consumes ATP/NADPH in the ratio 1.5, however linear electron transfer (LET) from water to NADP^+^ via photosystem II (PSII), plastoquinone/ plastoquinol (PQ/PQH_2_), cytochrome *b*_6_*f* (cyt*b*_6_*f*), plastocyanin (PC), photosystem I (PSI), ferredoxin (FD) and ferredoxin-NADP^+^-reductase (FNR) only produces 1.28 ATP/NADPH (Allen, 2002). The ATP/NADPH demands of the stroma can be further increased by the occurrence of photorespiration, the reaction of O_2_ rather than CO_2_ with ribulose bisphosphate (RuBP), catalysed by the CBB cycle enzyme Rubisco. The oxygenation reaction produces 2-phosphoglycolate, which then requires additional ATP to regenerate into 3-phosphoglycerate for the CBB cycle. In addition to photorespiration, a range of other biosynthetic processes including nitrogen fixation and protein and lipid biosynthesis in the chloroplast can also affect ATP/NADPH demand (Busch, 2020). Moreover, the balance between the production of ATP and NADPH by the light reactions and their consumption by the downstream metabolism can be further disturbed by the changing environmental conditions experienced by land plants, such as light intensity, temperature and water availability, which also impacts CO_2_ availability through stomatal closure (Kramer and Evans, 2010). The imbalances created can limit CO_2_ fixation and cause photoinhibition via the over-reduction of the chloroplast stroma and production of reactive oxygen species (ROS) (Li et al., 2009). Optimising how photosynthesis responds to these environmental disturbances has therefore been identified as a promising target to improve plant productivity (Wang et al., 2020; Long et al., 2022).

Imbalances in the production and consumption of ATP/ NADPH are most readily observed through the instability they create in CO_2_ assimilation rates, which are logically mirrored by similar instability in the chlorophyll fluorescence and absorption signals associated with proton (electrochromic shift, ECS) and PSI electron (P700) transport reactions during photosynthesis. These photosynthetic oscillations have been observed in a range of species including including broad bean, barley, tobacco, tomato, soybean, spinach and sunflower, and could be triggered by sudden shifts in light intensity, CO_2_ concentration, temperature or short dark periods (Sivak and Walker, 1983, 1986; Sivak et al., 1985; Stitt, 1986; Nakamoto et al., 1987; Stitt and Schreiber, 1988; McClain and Sharkey, 2023). The observed sequence of events during these oscillations suggests that transient ATP deficit in the chloroplast triggers a slowdown in NADPH consumption and thus over-reduction of the electron transport chain. Consistent with this, photosynthetic oscillations could also be exacerbated by feeding leaves with mannose or 2-deoxyglucose, which sequester inorganic Pi, which together with ADP and proton motive force (pmf) are required for ATP synthesis (Walker, 1992). Similarly, oscillations were demonstrated in wild-type tobacco plants via rapid ramping of CO_2_ concentration at 18°C, though interestingly the effect was absent at 23°C (McClain and Sharkey, 2023). The suggested cause was a failure to utilise the product of the CBB cycle triose phosphate due to a temperature limitation of sucrose synthesis, which normally regenerates Pi for the chloroplast (McClain and Sharkey, 2023).

To avoid photosynthetic oscillations and the associated losses in CO_2_ fixation, mechanisms must be present which dampen the imbalances by flexibly providing additional ATP. ATP level can be augmented via several alternative electron transfer pathways that create ATP without net production of NADPH. These include consumption of NADPH to reduce malate, which is exported to the mitochondria to power respiratory electron transfer (the malate-valve), the reduction of O_2_ by PSI (water-water cycle) or cyclic electron transfer (CET) (Miyake, 2010; Alric and Johnson, 2017; Chadee et al., 2021). CET recycles electrons from FD to the PQ pool via two separate pathways; the antimycin A-sensitive Proton Gradient Regulation 5 (PGR5)-dependent and NADH-dehydrogenase-like dependent (NDH) CET (Yamori and Shikanai, 2015). In C3 plants the PGR5 pathway plays a dominant role in pmf formation, which protects PSI and PSII via induction of non-photochemical quenching (NPQ) and photosynthetic control (PCON), respectively (Munekage et al., 2004; Suorsa et al., 2012; Wang et al., 2015; Strand et al., 2017b). However, under low light and in C4 plants the NDH pathway plays a more substantial role (Yamori et al., 2015; Munekage and Taniguchi, 2016; Ogawa et al., 2023). Moreover, mutant plants with increased ATP demand due to a disturbed CBB cycle, showed a high CET phenotype dependent on the NDH pathway (Livingston et al., 2010a,b; Strand et al., 2017b). On the other hand, the *hope2* mutant, which has increased ATP synthase conductivity due to a mutation in the gamma subunit of the complex, showed a high CET phenotype dependent on the PGR5 pathway (Degen et al., 2023).

Consistent with a key role for CET, incubation of isolated barley protoplasts with the PGR5-dependent CET inhibitor antimycin A induced oscillations upon a shift from low to high light intensity (Furbank and Horton, 1987). However, since antimycin A also targets mitochondrial complex III and so affects the respiratory electron transport chain, no unambiguous evidence for the crucial involvement of CET in suppressing photosynthetic oscillations exists. Moreover, it remains unclear whether photosynthetic oscillations are damped primarily via PCON to avoid over-reduction of PSI acceptors or via provision of additional ATP. To this end, we utilised the Arabidopsis *pgr5*, *ndho* and *hope2* mutants to show that PGR5 is specifically required to prevent photosynthetic oscillations and that this action does not depend on its ability to induce PCON. These results shed new light on the crucial role PGR5-dependent CET plays during photosynthesis and provide support for its direct role in ATP provision.

## Materials and Methods

### Plant material, growth conditions and generation of double cross

*Arabidopsis thaliana* WT Col-0 and mutants *pgr5*, *ndho* and *hope2* were grown in a controlled-environment chamber for at least 6 weeks at 21 °C /15 °C day/night, 60% relative humidity with an 8-h photoperiod at a light intensity of 200 µmol photons m^−2^ s^−1^. Double mutants were generated by crossing *pgr5* with *ndho*. Seeds from successful crosses were sown and allowed to self-fertilise. Seedlings of the cross were screened for diminished capacity to induce NPQ during fluctuating light using an Imaging PAM (Heinz Walz GmbH, Effeltrich, Germany). Successful crosses were then confirmed via Western blot using the PGR5 and NdhS antibodies (Agrisera, Sweden).

### Chlorophyll fluorescence and P700, PC and Fd absorption spectroscopy

The Dual-KLAS-NIR photosynthesis analyser (Heinz Walz GmbH, Effeltrich, Germany) was used for pulse-amplitude modulation chlorophyll fluorescence measurements and P700 absorption spectroscopy in the near-infrared (Klughammer and Schreiber, 2016; Schreiber and Klughammer, 2016). After plants had dark-adapted for at least 1 h and for each genotype, one leaf was used to generate differential model plots according to manufacturer’s protocol, which were used for online deconvolution to determine redox changes of P700, PC and Fd. Prior to each measurement, maximum oxidation of P700 was determined by using the preprogrammed NIRmax routine (see Degen *et al.,* 2023 for further details). Dark-fluorescence (F0) and maximal fluorescence (Fm) were determined prior to the onset of actinic light.

### Electrochromic shift measurements

Electrochromic shift was measured using a Dual-PAM analyzer with a P515/535 emitter/detector module (Heinz Walz GmbH, Effeltrich, Germany) (Schreiber and Klughammer, 2008; Klughammer et al., 2013). Plants were dark-adapted for at least 1 h prior to measurements. The raw ECS traces were normalised to the height of a 50 µs single turnover flash applied prior to onset of actinic light (see Degen *et al*., 2023 for further details).

### Gas-exchange measurements

Chlorophyll fluorescence and P515 measurements at different atmospheric CO_2_ concentrations were performed using a Licor 6400XT (LI-COR Biosciences, Lincoln, NE, USA) fitted with the 6400-15 extended reach 1 cm chamber, which allowed for control of reference CO_2_ concentration and was used in combination with the Dual-PAM and Dual-KLAS (as described above).

Measurements using the Dynamic Assimilation Technique^TM^ were performed using a Licor 6800, following the instructions in (Saathoff and Welles, 2021) and the manual for the Licor 6800 Bluestem OS v2.1 (p. 9-73 - 9-106). Reference CO_2_ was ramped from 50 ppm to 1500 ppm at various speeds. For steady-state ACi curves, a pre-programmed routine was used, with maximum wait times of 90-180 s for each CO_2_ concentration, following the suggestion in (Sharkey, 2019) using reference CO_2_ concentrations 50, 100, 200, 300, 350, 400, 450, 500, 550, 600, 700, 800, 1000, 1200, 1500 and 420 ppm at a saturating light intensity of 1000 µmol m^−2^ s^−1^. Prior to measurements, leaves were acclimated at 400 ppm reference CO_2_ at either 18 or 23 °C.

## Results

### Short dark pulses induce oscillations in *pgr5*

Chlorophyll fluorescence was measured in dark-adapted WT Col-0 and CET-deficient *pgr5* and *ndho* mutants over a 6 min actinic light period of 169 μmol photons m^−2^ s^−1^, which was interrupted periodically by a series of dark pulses (Fig.1). During the first 30 s, chlorophyll fluorescence (Chl F) responded similarly in the Col-0, *pgr5* and *ndho* mutants, and the level of fluorescence quenching was similar. However, in the subsequent minutes the fluorescence level in the WT and *ndho* mutants, declined more quickly and to a lower amplitude than *pgr5*, which rose slightly in the 60-180 second window before very slowly falling again. These differences are consistent with the low NPQ phenotype previously observed in *pgr5* (Supplemental Fig. S1) and the fact that *ndho* plants show no decrease and even sometimes a slight increase in NPQ (Munekage et al., 2002; Rumeau et al., 2005; Degen et al., 2023). In Col-0, the dark pulses resulted in a steep drop in Chl F and a slight rise after light was subsequently reapplied (Fig. 1a), although returned to steady state levels after 10 s. In contrast, after 120 s illumination and 5 dark pulses, Chl F in *pgr5* began to oscillate after each subsequent dark pulse (Fig. 1c). However, the NDH-deficient *ndho* mutant did not exhibit this extreme oscillatory behaviour after the dark pulses and behaved similarly to Col-0 (Fig. 1e). Previous experiments also observed oscillations in the ECS signal (Sivak et al., 1985), which provides information of the pmf amplitude, membrane proton conductivity and flux and the ATP/ADP ratio in the chloroplast (Joliot and Delosme, 1974; Kramer and Crofts, 1989; Sacksteder and Kramer, 2000; Baker et al., 2007; Buchert et al., 2021). In Col-0, the ECS signal behaved similarly to Chl F, only showing a slight increase after the 1.5 s dark pulse, but thereafter returned quickly to a steady-state level (Fig. 1b). In *pgr5* on the other hand, oscillations occurred after 120s in the light following each dark pulse, while in *ndho* (Fig, 1d, f) no oscillations were observed in the ECS signal which behaved like Col-0. This demonstrates that the oscillatory behaviour observed in the chlorophyll fluorescence signal of *pgr5* also affects its pmf.

**Figure 1:**
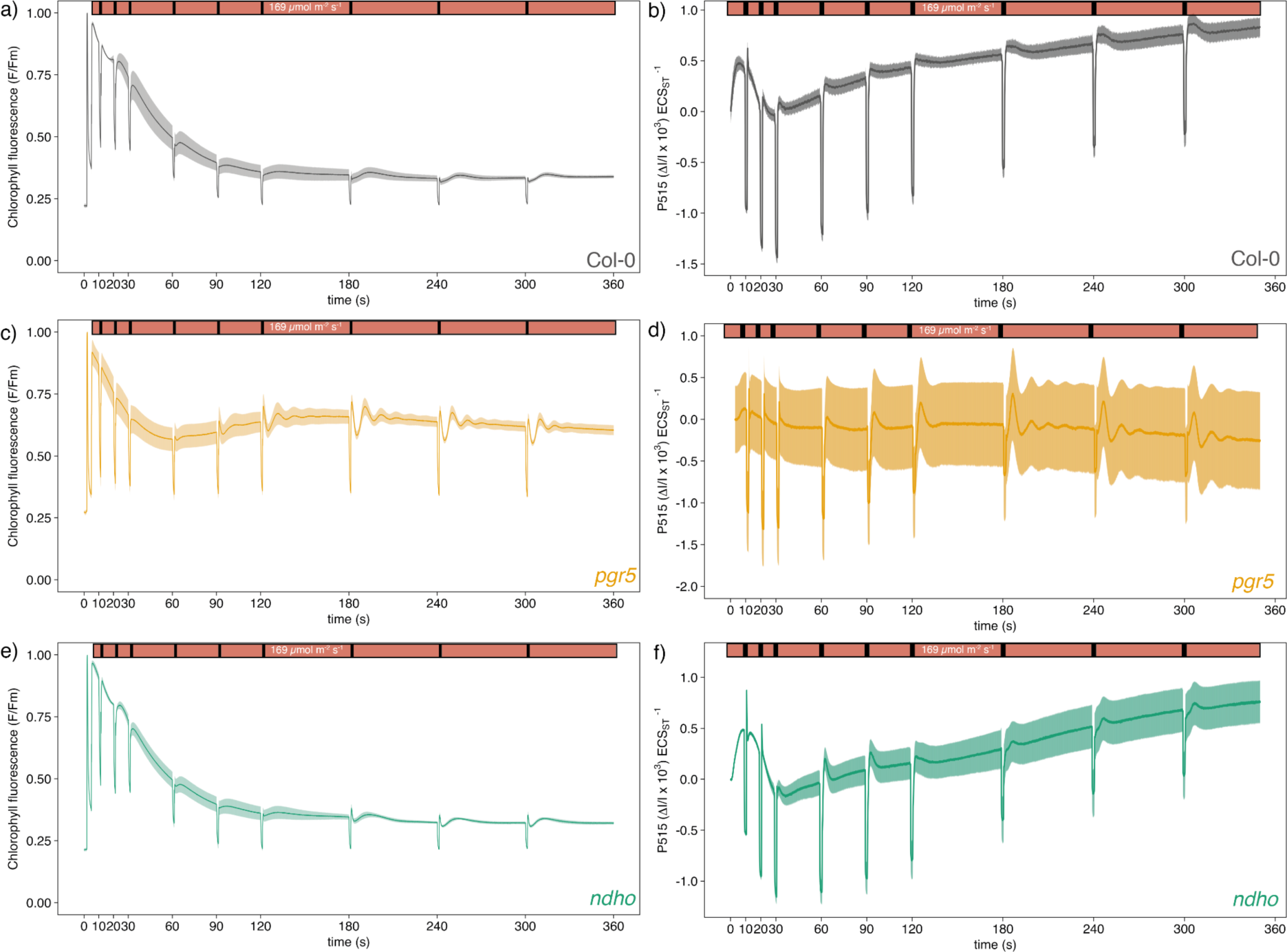
Chlorophyll fluorescence and electrochromic shift during photosynthetic induction. For chlorophyll fluorescence (a, c, e) Fo/Fm was measured before actinic light was turned on and fluorescence normalised to Fm. The electrochromic shift at 515 nm was normalised by the height of a 50 µs saturating pulse applied prior to the measurements. Dark pulses were applied at the times indicated. (a, b) Col-0, (c, d) *pgr5* and (e, f) *ndho*. Lines show the average of at least three biological replicates and shaded areas represent SEM.

Intriguingly, the above data shows that in *pgr5*, oscillations only occur after a certain period in the light, after 2 min they are observable in the Chl F trace and after 3 min in the ECS trace. Hence, we sought to establish whether preceding dark pulses or time in the light influence oscillations. Therefore, dark pulses were applied after 90 s, 180 s and 300 s (Fig. 2a). The dark pulse after 90 s did not induce oscillations in *pgr5*, whereas after 180 s and 300 s oscillations were induced. This confirms that only once the steady-state is reached in *pgr5* after 120 s (Fig. 2b), disruption of continuous illumination by short dark pulses results in oscillations.

**Figure 2:**
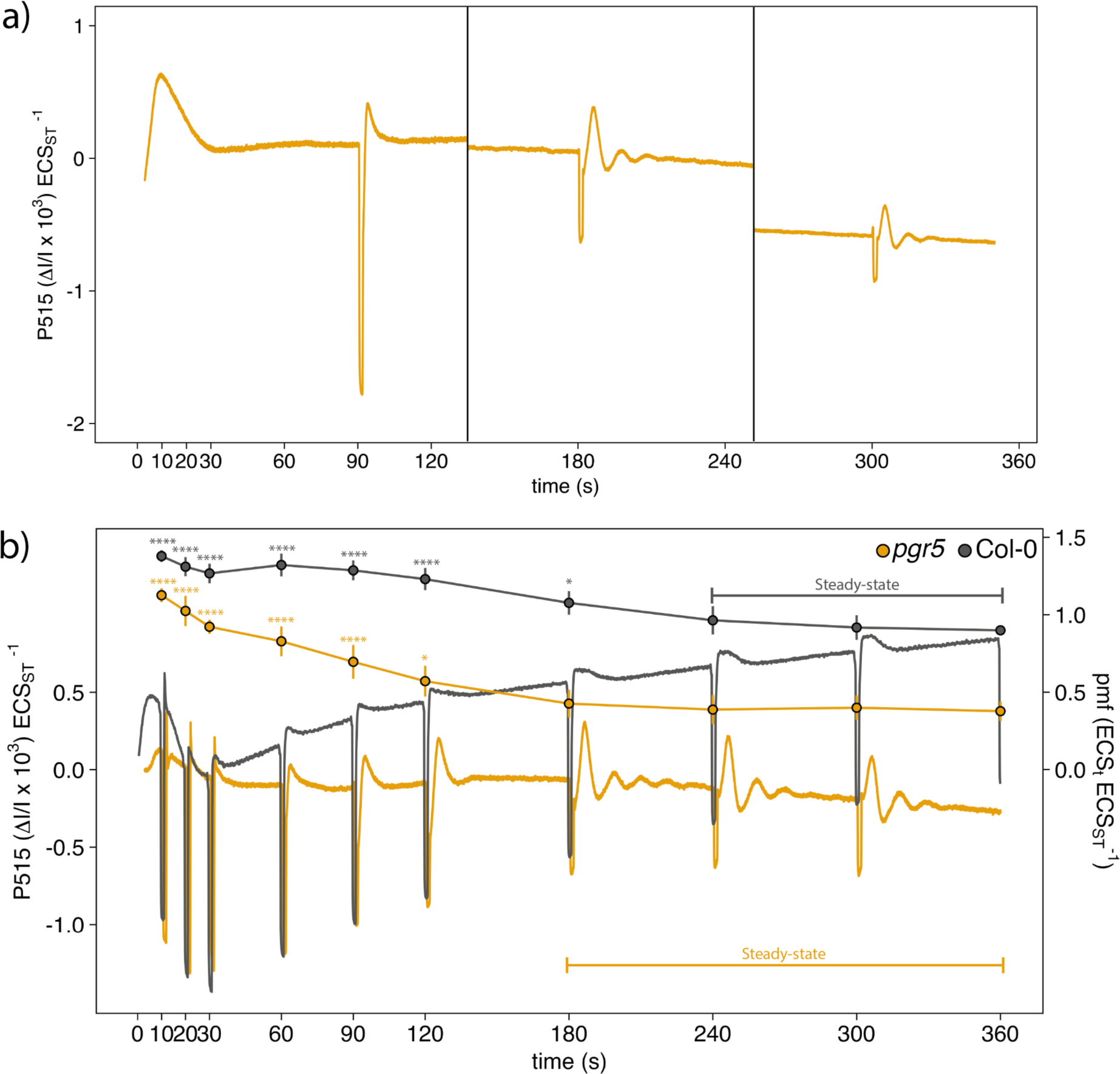
Oscillations in *pgr5* during steady-state. a) Short dark-pulses were applied at 90, 180 and 300s (separate measurements). b) pmf measurements of Col-0 and *pgr5* are plotted alongside raw ECS traces from Fig. 1. Asterisks indicate significant difference relative to the pmf value at 360s (2-way ANOVA). Steady-state was reached when pmf was not significantly different. Lines and points show the average of at least three biological replicates and SEM was omitted for better visual representation. Error bars represent SD.

### Oscillations can be induced in *pgr5* by specific changes in light intensity

Past work was able to induce oscillations by changing the light intensity rapidly (Furbank and Horton, 1987; Laisk et al., 1991). In Col-0, no oscillatory behaviour was observed when light intensity was increased from 59 µmol photons m^−2^ s^−1^ (Low Light (LL)) to 228 µmol photons m^−2^ s^−1^ (Medium Light (ML)) and to 628 µmol photons m^−2^ s^−1^ (High Light (HL)) (Fig. 3a). Whereas in *pgr5* the LL to ML change induced oscillations (Fig. 3c). Increasing light from ML to HL, however, did not result in oscillatory behaviour in *pgr5*. In *ndho* increasing light intensities did not result in oscillations, similar to Col-0 (Fig. 3e). Once again, the ECS signal was only found to oscillate in *pgr5* (Figs. 3b, d, f).

**Figure 3:**
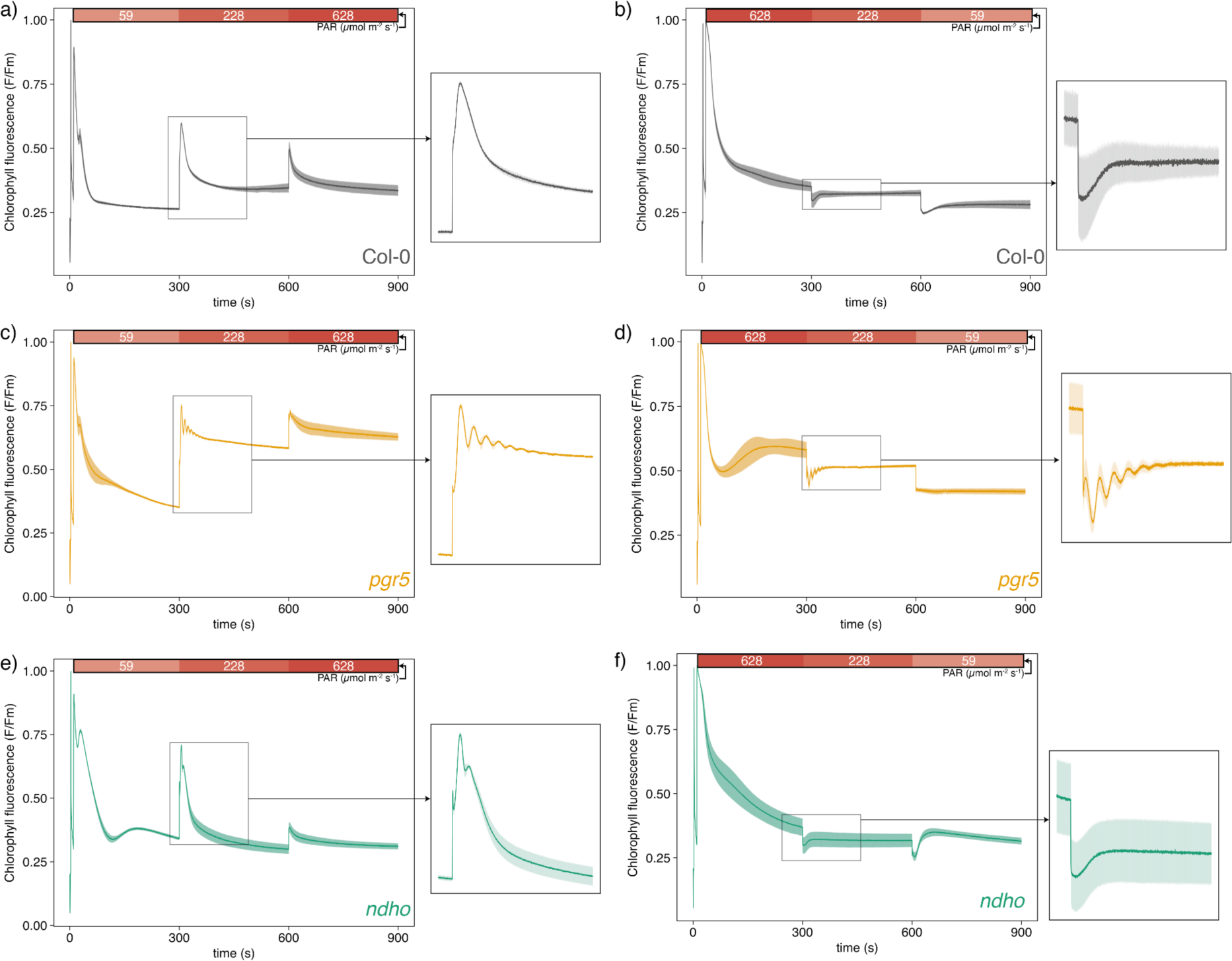
Chlorophyll fluorescence during increasing light intensity. a) Col-0, c) *pgr5*, e) *ndho*. Light intensity is indicated at the top of the plot. Fluorescence was normalised to Fm recorded previous to the measurements. P515 absorption changes during increasing light intensity. The P515 signal was normalised to the height of a 50 µs single turnover flash applied prior to measurements. b) Col-0, d) *pgr5*, f) *ndho*. Light intensity is indicated at the top of the plot. Lines show the average of at least three biological replicates and shaded areas represent SEM.

To determine whether the change in light intensity itself or the directionality of change was important to induce oscillations, we reversed the light intensity change from HL to LL. In Col-0 and *ndho* no oscillations were observed (Fig. 4a, c, e). In *pgr5*, the change from HL to ML light resulted in oscillations, however the change from ML to LL did not (Fig. 3d). This suggests that rather than the directionality of light intensity change, the extent of change in light intensity itself is important for observation of oscillations. As above a similar pattern in the ECS signal was observed (Fig. 4b, e, f), where only *pgr5* oscillated. Intriguingly, no oscillations were observed in *pgr5* if the light intensity was changed directly from low to high (Supplemental Fig. S2).

**Figure 4.**
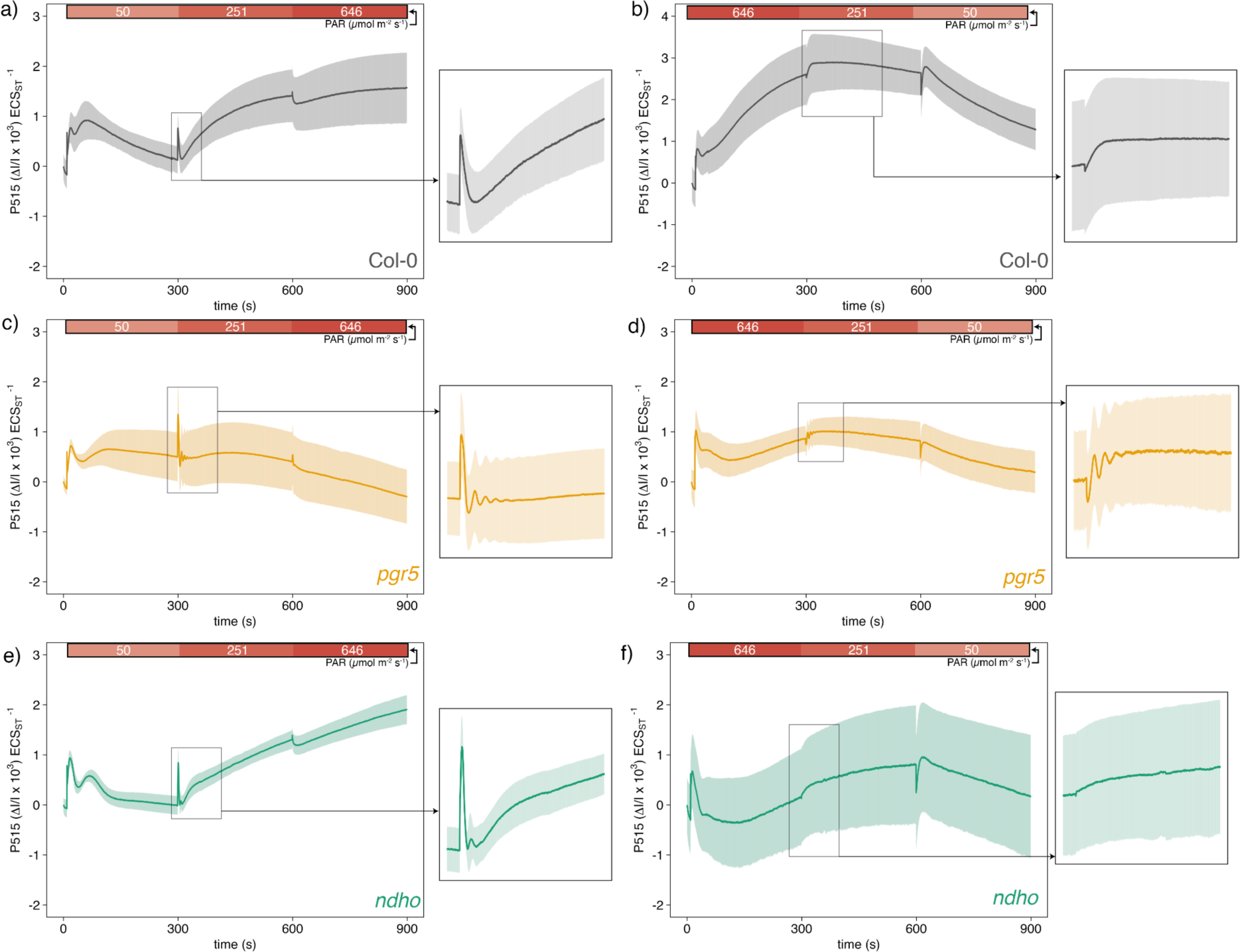
Chlorophyll fluorescence during decreasing light intensity. a) Col-0, c) *pgr5*, e) *ndho*. Light intensity is indicated at the top of the plot. Fluorescence was normalised to Fm recorded previous to the measurements. P515 absorption changes during decreasing light intensity. The P515 signal was normalised to the height of a 50 µs single turnover flash applied prior to measurements. b) Col-0, d) *pgr5*, f) *ndho*. Light intensity is indicated at the top of the plot. Lines show the average of at least three biological replicates and shaded areas represent SEM.

### Oscillations in *pgr5* are dampened at low and high CO_2_

A shift from ambient air to high CO_2_ has previously been shown to induce oscillations (Walker et al., 1983). To this end, we exposed leaves to varying CO_2_ concentrations for 5 min and then suddenly changed the light intensity from 190 µmol photons m^−2^ s^−1^ to 646 µmol photons m^−2^ s^−1^. In Col-0, this resulted in an increase in chlorophyll fluorescence for ca. 5s, which was subsequently quenched (Fig. 5a). The amplitude of fluorescence increase was highest at 400 ppm and somewhat lower at 200 and 100 ppm, while the subsequent rate of quenching was faster in the order 100 > 200 > 400 ppm. At 800 and 1200 ppm, the fluorescence rise was smaller and the subsequent quenching much slower. We observed the ECS rose more sharply following the light shift at 100 ppm consistent with the higher pmf implied by the more rapid quenching of Chl F (Fig. 5c). Whereas, at 800 and 1200 ppm the ECS change was minimal suggesting the higher CO_2_ concentration mitigated the rise in pmf in line with a larger sink capacity.

**Figure 5:**
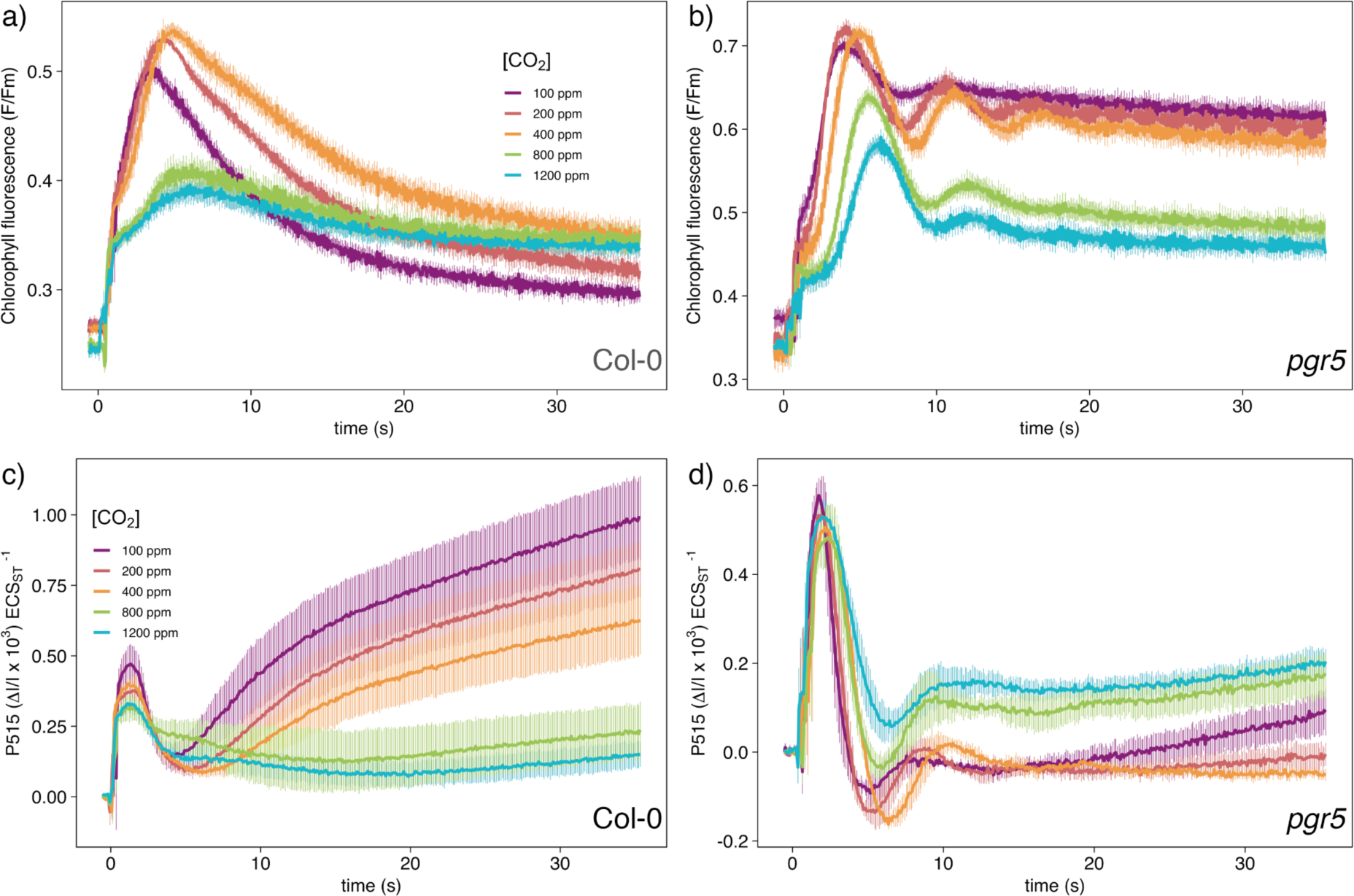
Chlorophyll fluorescence and P515 signals in response to CO_2_ concentrations. a,c) Col-0, b,d) *pgr5*. At time point 0 actinic light was increased from 190 µmol photons m^−2^ s^−1^ to 646 µmol photons m^−2^ s^−1^. Leaves were acclimated for 5 min at each CO_2_ concentration. Lines show the average of at least three biological replicates and error bars represent SEM.

The picture in *pgr5* was quite different with oscillations induced upon an increase in light intensity at each CO_2_ concentration (Fig. 5b). The oscillations were most intense at 400 and 200 ppm, although were somewhat dampened at 100 ppm. In each case the final fluorescence level was much higher than in Col-0. The light intensity shift combined with 800 and 1200 ppm CO_2_ also dampened the oscillations in *pgr5* and resulted in a lower final fluorescence level (Fig. 5b). We observed a similar behaviour in the ECS signal (Fig. 5d), although oscillations in *pgr5* were less prominent than in the Chl F signal, though still dampened at 100 ppm and 800 and 1200 ppm CO_2_. Noticeably, the ECS signal was higher in *pgr5* at 800 and 1200 ppm following the light shift compared to 100-400 ppm suggesting that under low CO_2_ pmf generation is limited in the mutant but that this is somewhat mitigated by higher sink capacity (Fig. 5b).

### Oscillations induced by the Dynamic Assimilation Technique^TM^ are a distinct phenomenon

Recently, McClain et al. (2023) observed that during much more rapid ramping of CO_2_ concentrations using the Dynamic Assimilation Technique^TM^ (DAT) (Saathoff and Welles, 2021), photosynthetic oscillations could be induced in tobacco. These conditions were able to trigger triose phosphate utilisation (TPU) limitation, i.e. an inhibition of photosynthesis caused by depleted chloroplast Pi concentration triggered by a limitation on the rate of TP conversion into sucrose in the cytoplasm at low temperature. To this end, we used DAT to verify whether oscillations of this type might also be linked to CET in *Arabidopsis*. Steady-state ACi curves of *pgr5* had diminished capacity for CO_2_-assimilation compared to Col-0 as reported previously (Munekage et al., 2008; Degen et al., 2023). This was the case at a leaf temperature of both 23 °C and 18 °C (Fig. 6 a,b). DAT with a ramp of 100 ppm/min resulted in higher A values in Col-0 at 23 °C and slightly lower at 18 °C compared to the steady state (Fig. 6a). However, in *pgr5* CO_2_-assimilation at 18 °C was higher and similar to the ACi curve of *pgr5* at 23 °C, suggesting the limitations at low temperature encountered by Col-0 may not apply in the mutant (Fig. 6b). At faster ramps of 400 ppm/min and 600 ppm/min, CO_2_-assimilation at 18 °C in *pgr5* was higher compared to 23 °C. However, no oscillations in CO_2_-assimilation were observed in *pgr5*. At 18 °C leaf temperature in Col-0, on the other hand, ramps at 400 ppm/min caused a distinct peak and trough in CO_2_ assimilation. At 600 ppm/min the ACi curve consisted of a single clearly defined peak. Collectively these data demonstrate that the oscillations induced by DAT in the WT are a different phenomenon from those induced by dark pulses, changing light intensity or CO_2_ concentration in *pgr5*. Transforming and plotting the data in Fig. 6c, d so that CO_2_-assimilation is plotted against the log10 of time makes it clear that a faster ramping speed resulted in a higher assimilation at 18 °C in *pgr5*. This was also the case for Col-0 at 18 °C, however, at 23 °C a ramp of 100 ppm/min resulted in higher than steady-state CO_2_ assimilation.

**Figure 6:**
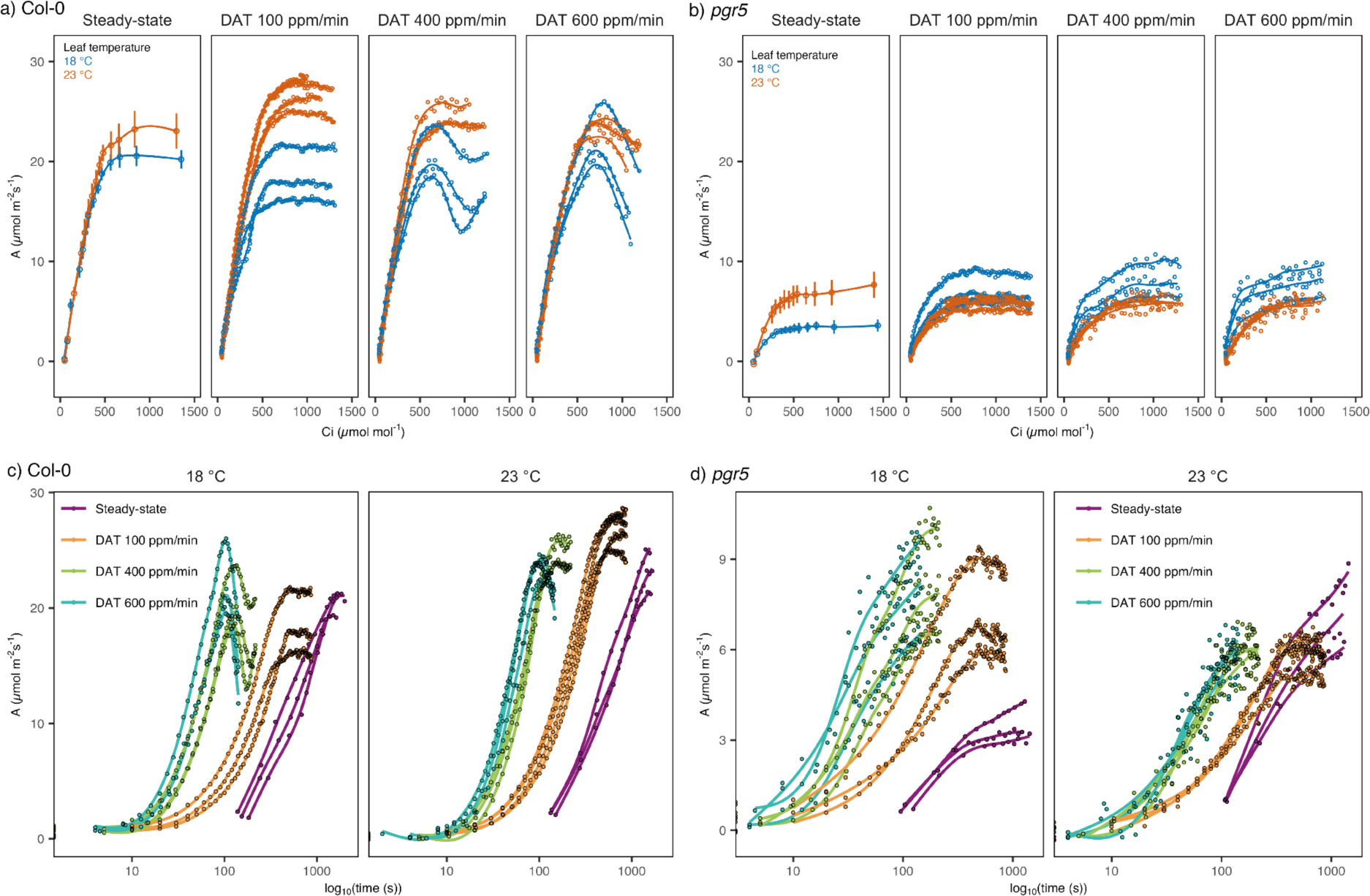
CO2-assimilation curves using the Dynamic Assimilation Technique ™ at various ramp rates. a,c) CO_2_-assimilation in Col-0, b,d) CO_2_-assimilation in *pgr5*. a,b) ACi curves using steady-state CO_2_ increases and DAT ramp speeds of 100,400 or 600 ppm/min. c,d) Data from A,B replotted on a log_10_ scale. Colours indicate different leaf temperatures or steady-state and DAT ramp speeds. For the steady-state curves in a and b, mean and standard deviation are plotted (n=3). For DAT curves in a-d, A/Ci curves of individual plants are shown, separated by a smoothed non-linear fit.

### Oscillations in *pgr5* are unrelated to the absence of photosynthetic control

Photosynthetic oscillations also involve transient reduction in PSI (P700) suggesting they trigger overeduction of the stromal NADPH pool (McClain and Sharkey, 2023). It was therefore interesting to examine the extent to which lack of PCON, which mitigates against P700 reduction by regulating electron flow between cyt*b*_6_*f* and PSI, would affect oscillations. To this end, we compared the *pgr5* with the *hope2* mutant, which has a mis-regulated ATP synthase and was shown to possess a similar low donor-side limitation of PSI (Y(ND)) and high acceptor-side limitation (Y(NA)) phenotype in addition to similar high proton conductivity (Degen et al., 2023). We found *hope2* did not exhibit any oscillatory behaviour upon application of short dark intervals, changes in light intensity or CO_2_ (Fig. 7 and Supplemental Fig. S2, S3), demonstrating that the absence of PCON is not the determining factor in the oscillations. In contrast to the similarities with *pgr5*, *hope2* is able to maintain a WT-like pmf amplitude by increasing PGR5-dependent CET (Degen et al., 2023). Therefore, our data implicates the low amplitude of the pmf in *pgr5* as the causative factor in provoking the oscillations. To confirm that the upregulated PGR5-dependent CET suppresses oscillations in the *hope2* background we examined the *hope2 pgr5* cross (Fig. 7c). Consistent with dependence of oscillations on the absence of PGR5, oscillations were restored in the double mutant, however the *hope2 ndho* double mutant was unaffected (Fig. 7c).

**Figure 7:**
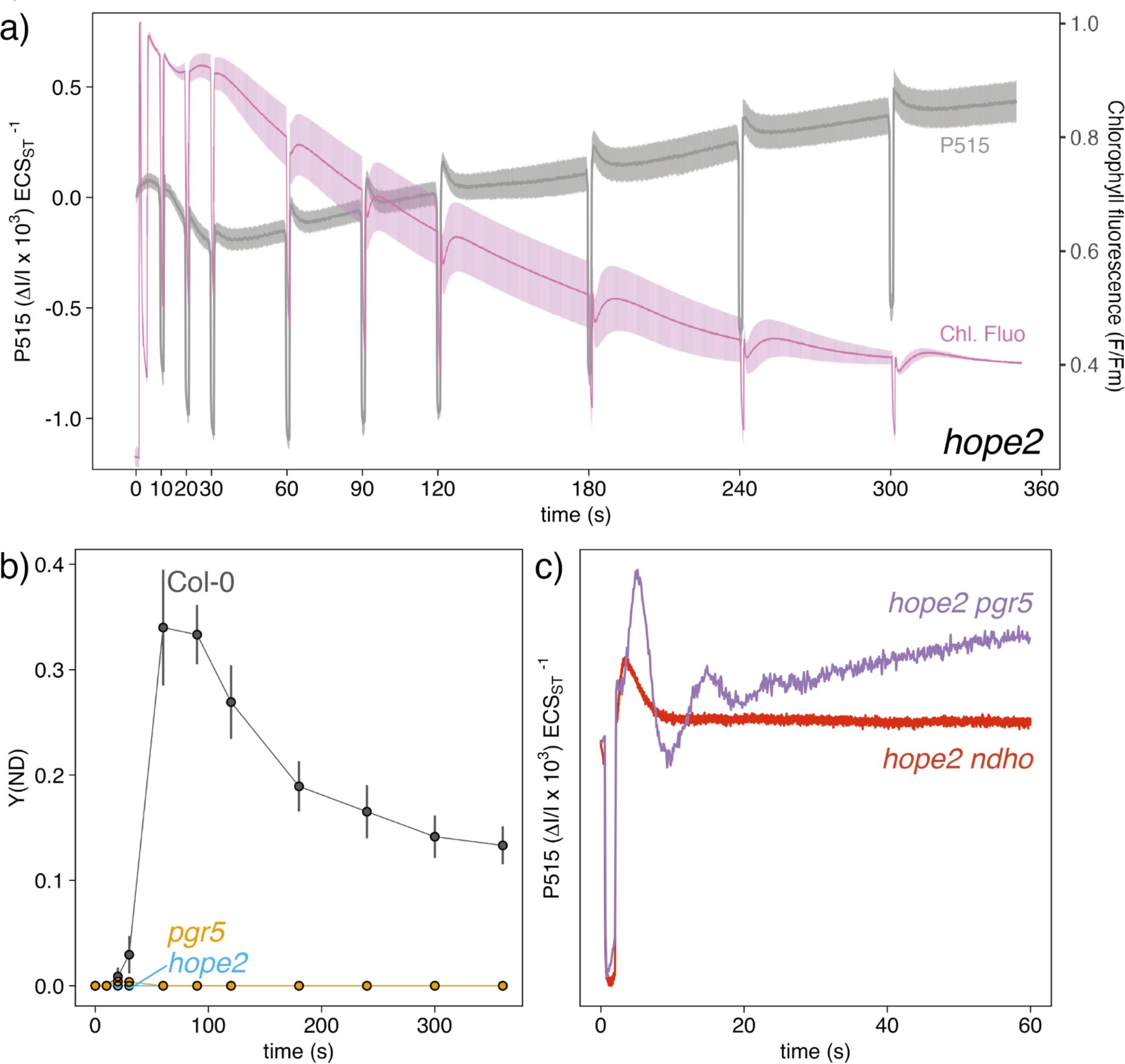
Photosynthetic characteristics of *hope2* and double mutants. a) Chlorophyll fluorescence trace, normalised to Fm and P515 absorption changes during photosynthetic induction. The P515 signal was normalised to the height of a 50 µs single turnover flash applied prior to measurements. b) Lack of photosynthetic control (Y(ND) in *hope2* and *pgr5* during photosynthetic induction c) Photosynthetic oscillations in the P515 signal after a short dark pulse are present in *hope2 pgr5* although not *hope2 ndho*. Colours indicate different signals or genotypes, where indicated. Lines show the average of at least three biological replicates and shaded areas represent SEM. Symbols in (b) represent the mean of three biological replicates +/−SD.

### Oscillations in *pgr5* depend on loss of PGR5 protein and are increased in severity by loss of the NDH complex

Since the original *pgr5* mutant was generated by EMS mutagenesis, we also checked for the presence of oscillations in the *pgr5*-CAS mutant (Penzler et al., 2022), which does not contain any other background mutations (Fig. 8). The data confirm that the oscillations are specifically associated with the lack of PGR5. Using the *pgr5*-CAS mutant we then determined the phase relationship between the oscillations observed within the various photosynthetic signals in *pgr5* including Chl F (which reports on the redox state of the PSII acceptor Q_A_^−^), ECS, PC, P700 and Fd (Fig. 9). In both Col-0 and *pgr5-CAS*, upon imposition of a dark pulse we observe the instantaneous oxidation of Fd as light-driven electron transport is halted, while pmf also drops rapidly. The two genotypes then follow different paths. In Col-0, as light is reimposed, reduced Fd and oxidised PC, P700 transiently accumulates with the formation of ∼50% of the total pmf. There is then a very brief lag phase (∼0.25 s) coinciding with transient re-reduction of PC and P700 and the re-oxidation of Fd. This re-oxidation of Fd in the light precedes the formation of the remaining 50% of the pmf and the complete oxidation of PC and P700. In *pgr5*, the same sequence of events observed in Col-0 follows the dark pulse, however, here the key difference is the much slower re-oxidation of Fd and a much longer accompanying lag-phase (∼1-2 s) in pmf formation (highlighted in Fig. 9e with a shaded box). This lag-phase coincides with a much more severe transient reduction in Q_A_^−^, PC and P700 confirming that the LET chain is over-reduced by the failure to re-oxidise Fd rapidly during this period.

**Figure 8:**
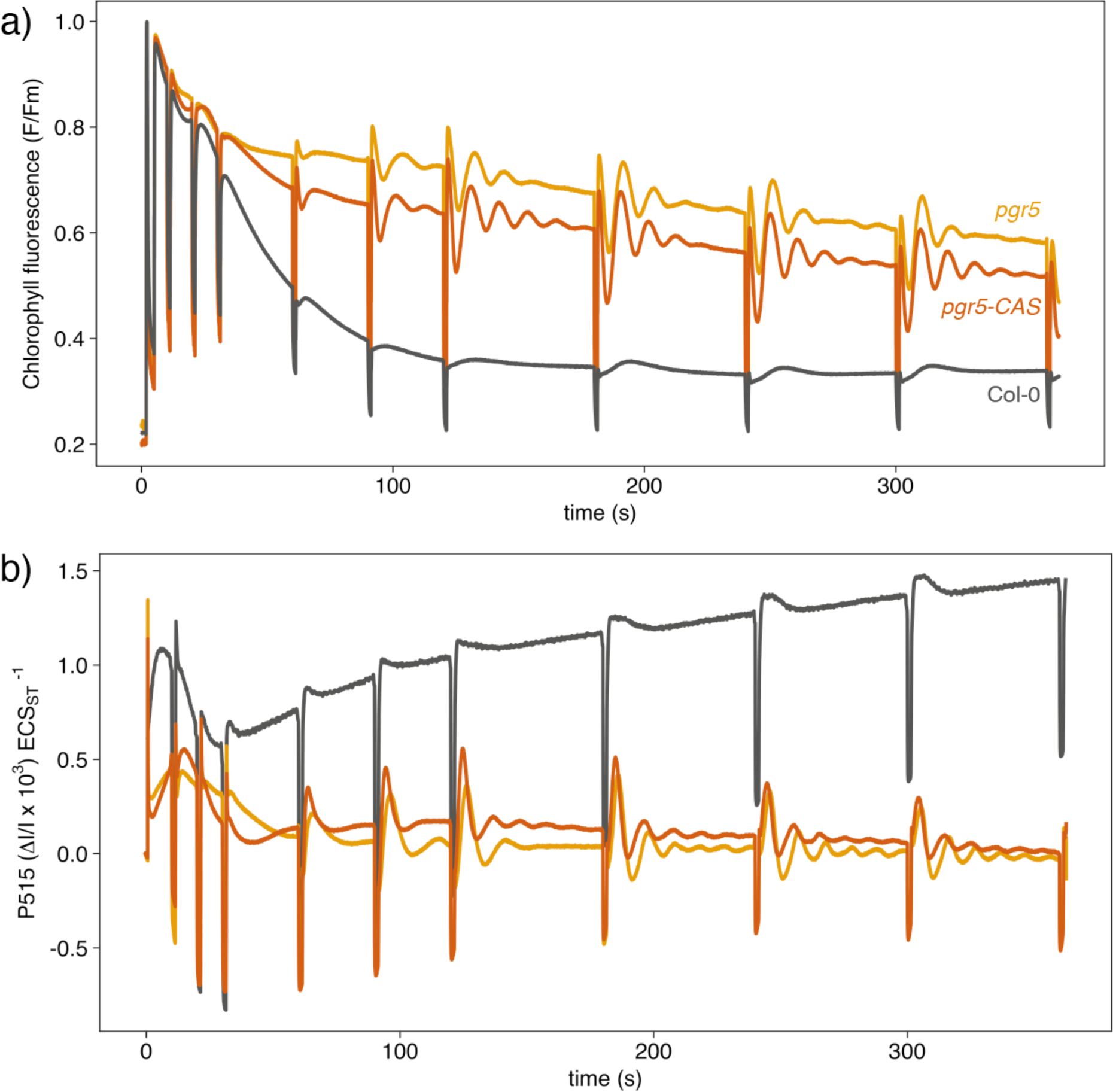
Oscillations in Col-0, *pgr5* and *pgr5*-CAS. a) Chlorophyll fluorescence, b) Electrochromic shift (P515). Red actinic light was interrupted by 1.4 s dark periods to induce oscillations. Fluorescence was normalised to the maximum value determined by a 300 ms flash. P515 was normalised to the height of a 50 µs single-turnover flash. Colours indicate different genotypes. Lines show the average of at least three biological replicates and SEM was omitted for better visual comparison.

**Figure 9:**
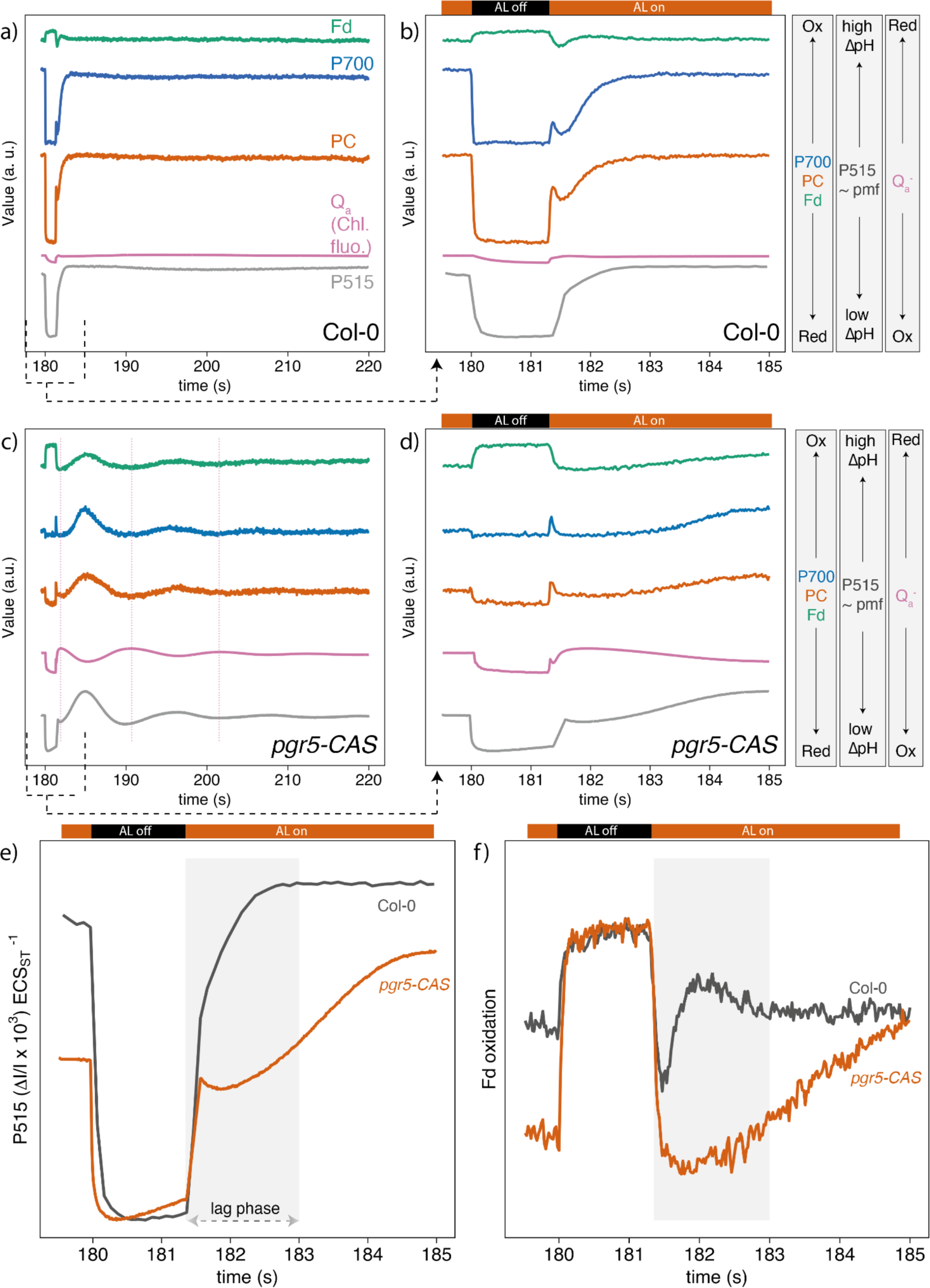
Phase-relationship between oscillatory components of the electron transport chain of Col-0 and *pgr5-CAS*. a,b) Raw traces of Col-0. Cc, d) Raw traces of *pgr5-CAS*. b, d) represent zoomed versions of a and c. Vertical dashed lines indicate peaks of chlorophyll fluorescence which anticipate peaks of other oscillatory components. e) Raw ECS trace after the short dark period. The grey box indicates the lag phase in pmf in *pgr5-CAS*. f) Raw traces of the Fd signal after the short dark period. Colours indicate oscillatory components or genotypes, where indicated. Lines show the average of at least three biological replicates and SEM was omitted for better visual comparison.

Interestingly, in the double mutant *pgr5 ndho* (Supplemental Fig. S4), the oscillations following a dark pulse were even more pronounced than in the single *pgr5* mutant. This demonstrates that while PGR5 can largely compensate for the loss of NDH, there is some compensation for the loss of PGR5 by NDH, confirming a cross talk between the two CET pathways in suppressing photosynthetic oscillations (Fig. 10).

**Figure 10:**
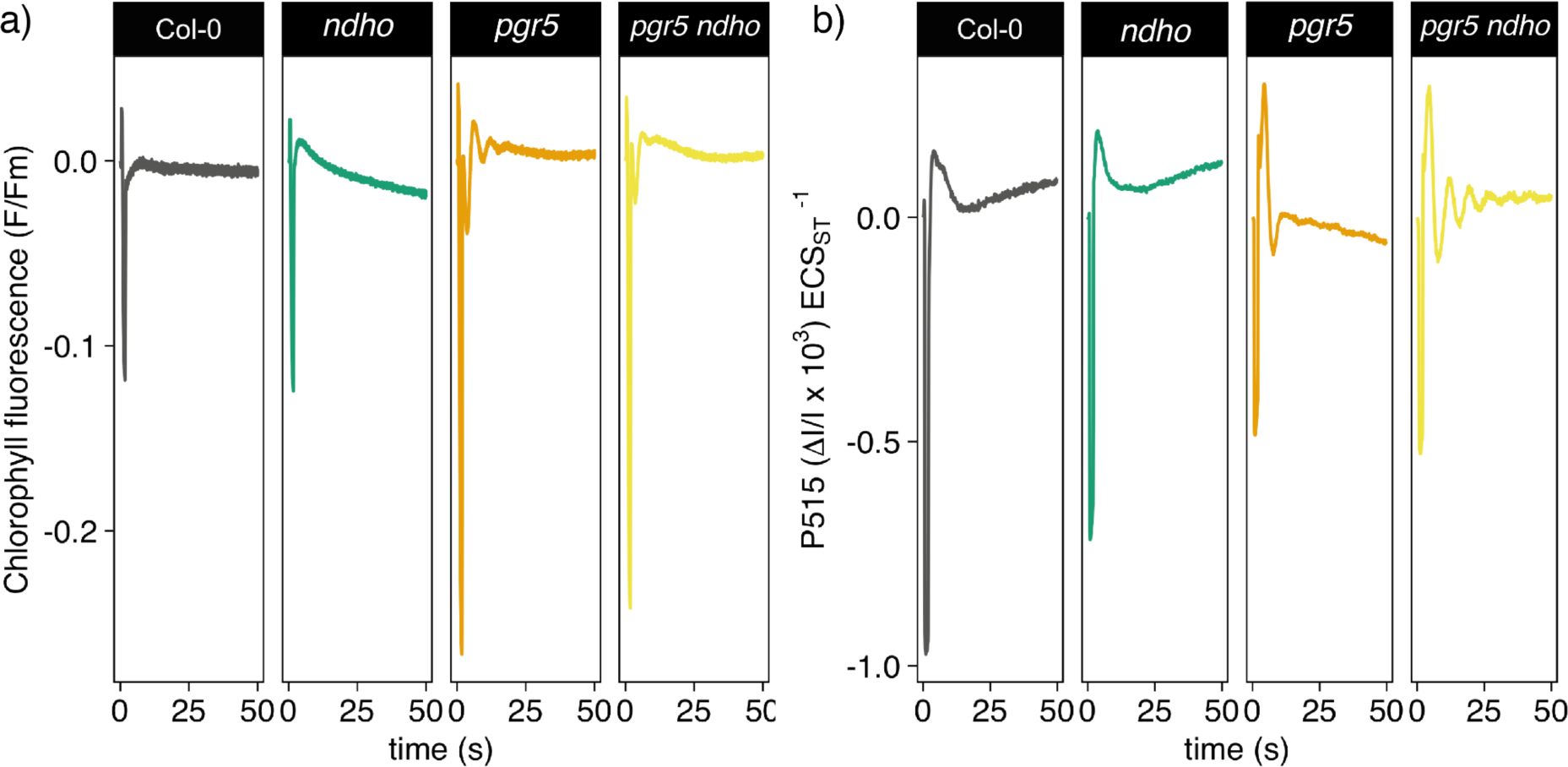
Photosynthetic oscillations in *pgr5 ndho*. a) Chlorophyll fluorescence and b) ECS signal. Red actinic light was interrupted by a brief dark pulse at time point 0. The P515 data was normalised to the height of 50 µs saturating single-turnover flash and chlorophyll fluorescence to Fm. Lines show the average of at least three biological replicates and SEM was omitted for better visual comparison. Colours indicate different genotypes.

## Discussion

### Independent of PCON, CET plays an important role in the suppression of photosynthetic oscillations

Photosynthetic oscillations lead to losses in CO_2_ fixation and the transient over-reduction of the electron transfer chain, which can in turn promote photo-oxidative stress. Interestingly however, despite net losses, photosynthesis during the oscillations can transiently outperform the steady-state rate (Sivak and Walker, 1986; Furbank and Horton, 1987; Laisk et al., 1991). Therefore, a more complete understanding of this phenomenon could engender future improvements in the ability of crops to withstand environmental shifts, while minimising carbon losses and photodamage. Oscillations have previously been explored in a range of C3 species including broad bean, barley, tobacco, tomato, soybean, spinach and sunflower (Ogawa, 1982; Sivak et al., 1985; Furbank and Horton, 1987; Stitt and Schreiber, 1988; Laisk et al., 1991; Stitt et al., 1991a,b). However, no oscillations were observed in the C4 species *Zea mays* and A*maranthus retroflexus*. The conclusion of this past work was that oscillations reflect imbalances in the ATP and NADPH levels required for optimal CBB cycle activity (Furbank and Horton, 1987; Laisk et al., 1991; Walker, 1992). This is corroborated by the observation that tobacco plants with decreased Rubisco content exhibited diminished oscillatory behaviour, likely due to diminished demand for ATP from the CBB cycle (Stitt et al., 1991b). Transitions in light intensity, temperature or CO_2_ concentration can induce oscillations, however, efforts to understand this phenomenon in greater detail have been hampered by the lack of available mutants which show enhanced photosynthetic oscillations. Given the central role advocated for CET in tuning the ATP/NADPH ratio to meet the energy and reducing power demands of the chloroplast, it is logical to suggest that this regulatory mechanism might play a key part in suppressing photosynthetic oscillations. Indeed, previous modelling and *in vitro* work was consistent with this idea (Furbank and Horton, 1987; Horton and Nicholson, 1987). Nonetheless, evidence that ATP/ NADPH levels are perturbed in mutants lacking CET, are scarce. ATP production in isolated *pgr5* chloroplasts was shown to be less efficient under CET conditions where Fd was provided as the sole electron acceptor, while *ndho* was unaffected (Wang et al., 2018). Another study found that ATP concentration as measured using the nanolantern fluorescent biosensor rose more slowly in *pgr5* and fell more rapidly (Sato et al., 2019). Alternatively, as shown in green algae *Chlamydomonas*, ATP levels in the *pgr5* mutant may be rescued largely through redundancy with other alternative electron transfer (AET) pathways, such as the water-water cycle and malate valve (Dang et al., 2014; Alric and Johnson, 2017). Indeed, much of the *pgr5* phenotype can be explained by the absence of PCON, suggesting the key role of CET may be in photoprotection of PSI rather than augmentation of ATP (Suorsa et al., 2012, 2016). However, absence of PCON does not explain the effects observed in our study since *hope2* lacks PCON and suffers high Y(NA) similar to *pgr5* (Degen et al., 2023), yet still failed to show oscillations. Moreover, regulation of pmf via the ATP synthase can also be ruled out since both *pgr5* and *hope2* show a similar high proton conductivity (Degen et al., 2023). Finally, the existence of some limited compensation of loss of PGR5 by the NDH pathway, as revealed in the double mutant *ndho pgr5* further implicates CET as playing a key role in the suppression of photosynthetic oscillations. The larger role we find for PGR5-dependent CET relative to NDH-dependent CET is consistent with its greater capacity in Arabidopsis (Munekage et al., 2004; Strand et al., 2017a). The key role for CET is also in line with the observation that oscillations were not observed in C4 plants, which have higher ATP demand and thus increased CET capacity (Ogawa, 1982; Nakamura et al., 2013; Munekage and Taniguchi, 2016; Ogawa et al., 2023).

### Oscillations in *pgr5* are dampened during photosynthetic induction and high and low CO_2_ concentrations

We found the oscillations in *pgr5* only appeared after dark pulses once the plants had been illuminated for ∼90-120 s in the light. This delay in the appearance of oscillations may be related to the higher pmf that is sustained in *pgr5* during induction (Fig. 2). This behaviour has been observed previously in *pgr5* and is related to the increase in pmf driven by restricted ATP synthase conductivity during induction, which relaxes as downstream electron sinks become active (Nikkanen et al., 2018; Degen et al., 2023). Consistent with this explanation, the ability of low CO_2_ to suppress the oscillations in *pgr5* is likely caused by the downregulation of ATP synthase conductivity under these conditions, independent of CET. In contrast, the partial suppression of oscillations by high CO_2_ in *pgr5* likely reflects the lowered ATP demand brought about by the suppression of photorespiration. This is in step with past work which showed the growth and fitness phenotype of *pgr5* is ameliorated by high CO_2_ (Munekage et al., 2002, 2008).

### Oscillations in *pgr5* do not depend on TPU limitation

In contrast with recent work by McClain *et al*., (2023) we can rule out that the oscillations in *pgr5* are caused by TPU limitation since they are not detected under rapid CO_2_ ramping unlike in the WT. This may reflect the fact that the *pgr5* mutant has a sufficiently low photosynthetic rate that it never enters TPU limitation. This conclusion is straightforwardly reconciled with the findings of McClain *et al*., 2023 that the overshoot in photosynthetic rate provoked by the sudden CO_2_ ramp unbalances metabolite pools in the CBB cycle. The effect of the ramp is to transiently utilise more Pi in ATP synthesis than can be regenerated by sucrose synthesis. This effectively produces a burst of photosynthesis followed by a trough as Pi is depleted and phosphorylation of 3-phosphoglycerate to 1,3-bisphosphoglycerate is slowed down. In turn this causes overeduction of the LET chain as NADPH consumption falls due to substrate depletion for glyceraldehyde dehydrogenase. We speculate in *pgr5* the lack of pmf precludes much of the rapid increase in photosynthetic rate upon CO_2_ ramp due to the existing lack of ATP. The cause of the faster rate of assimilation at lower temperatures in *pgr5* is puzzling and requires further experimentation to fully understand. It is possible that inhibition of sucrose synthesis at low temperature depletes Pi, raising pmf sufficiently in *pgr5* that PCON can operate. In this scenario the increase in the photosynthetic rate in *pgr5* may reflect better poising of the electron transfer chain, thus allowing more efficient operation of the NDH-dependent CET pathway and increased ATP production.

### Optimal suppression of oscillations requires rapid Fd oxidation and pmf generation by PGR5-dependent CET

Having ruled out the involvement of TPU limitations in inducing oscillations in *pgr5*, we looked to our simultaneous measurements of Fd, PC, P700, PSII fluorescence and ECS signals in whole leaves to understand the phase relationship and so cause and effect involved in photosynthetic oscillations. We observed that the reduction of P700 and Q_A_^−^ are out of phase with the changes in pmf in *pgr5*. Upon imposition of a dark pulse we observe the instantaneous oxidation of Fd as light-driven electron transport is halted, while pmf also drops rapidly. In contrast, as light is reimposed, reduced Fd accumulates more rapidly than pmf, resulting in a transient overreduction of the LET chain (as seen in P700, PC and Q_A_^−^ (Chl F) signals). Only upon the slow secondary rise of the pmf in *pgr5* do P700, Pc and Fd and Q_A_^−^ begin to become oxidised. In contrast in the WT, the secondary rise in the pmf is much more rapid and this suppresses the oscillations that ensue in the *pgr5* mutant. This lag phase in *pgr5* may reflect the time required to activate alternative mechanisms of pmf and ATP generation such as NDH-dependent CET or the water-water cycle. The subsequent photosynthetic oscillations that then ensue in *pgr5* may be caused by ‘over-oxidation’ of Fd as LET is reactivated by the sudden surge of ATP provided by the pmf rise. Normally in the WT, balance must be struck between two extremes, sufficiently oxidised Fd to allow LET *and* sufficiently reduced Fd to allow CET/AET to augment ATP production. We hypothesise based on these data that in the *pgr5* mutant this optimisation point is shifted because the NDH and water-water cycle may require a more reduced Fd pool to operate efficiently than does the PGR5 pathway. Thus, the *pgr5* mutant is crippled by oscillation between over-oxidised and over-reduced extremes, limiting carbon fixation following an environmental shift and thus increases the potential for photo-oxidative damage. The key attribute of the PGR5-dependent CET therefore may be its ability to respond rapidly to transient overreduction generating additional pmf and so ATP without becoming subsequently inhibited by the result of its own oxidative action on Fd.

## Author contributions

GED and MPJ designed research. GED performed experiments with help from FP. GED analysed data. GED and MPJ wrote the manuscript. All authors approved of the manuscript prior to submission.

---

**Supplemental Figure S1:**
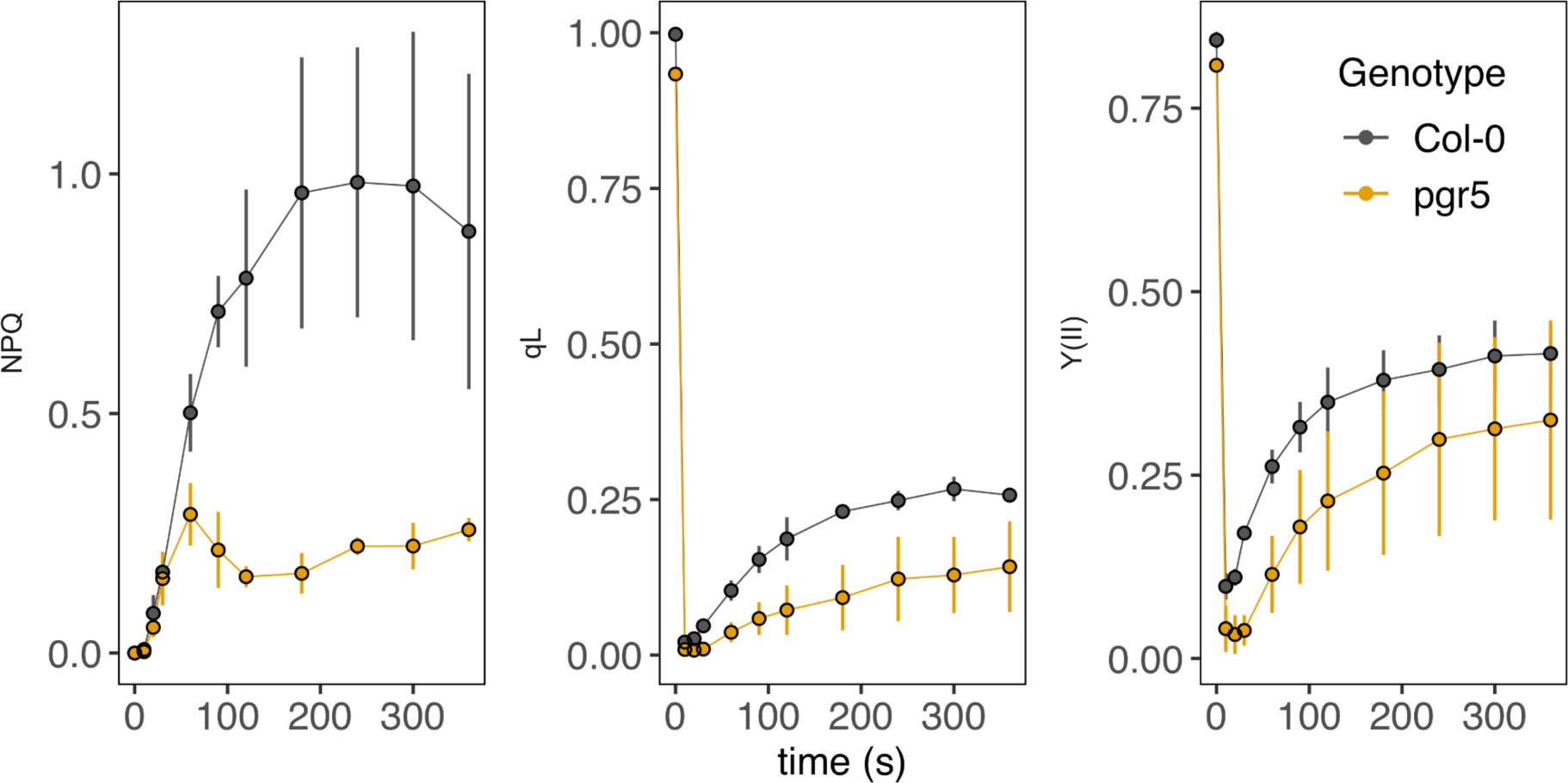
Photosynthetic parameters during induction. At time 0 Fm was determined with a saturating flash, following onset of actinic light. Symbols represent the mean of three biological replicates +/− SD. Colours indicate different genotypes.

**Supplemental Figure S2:**
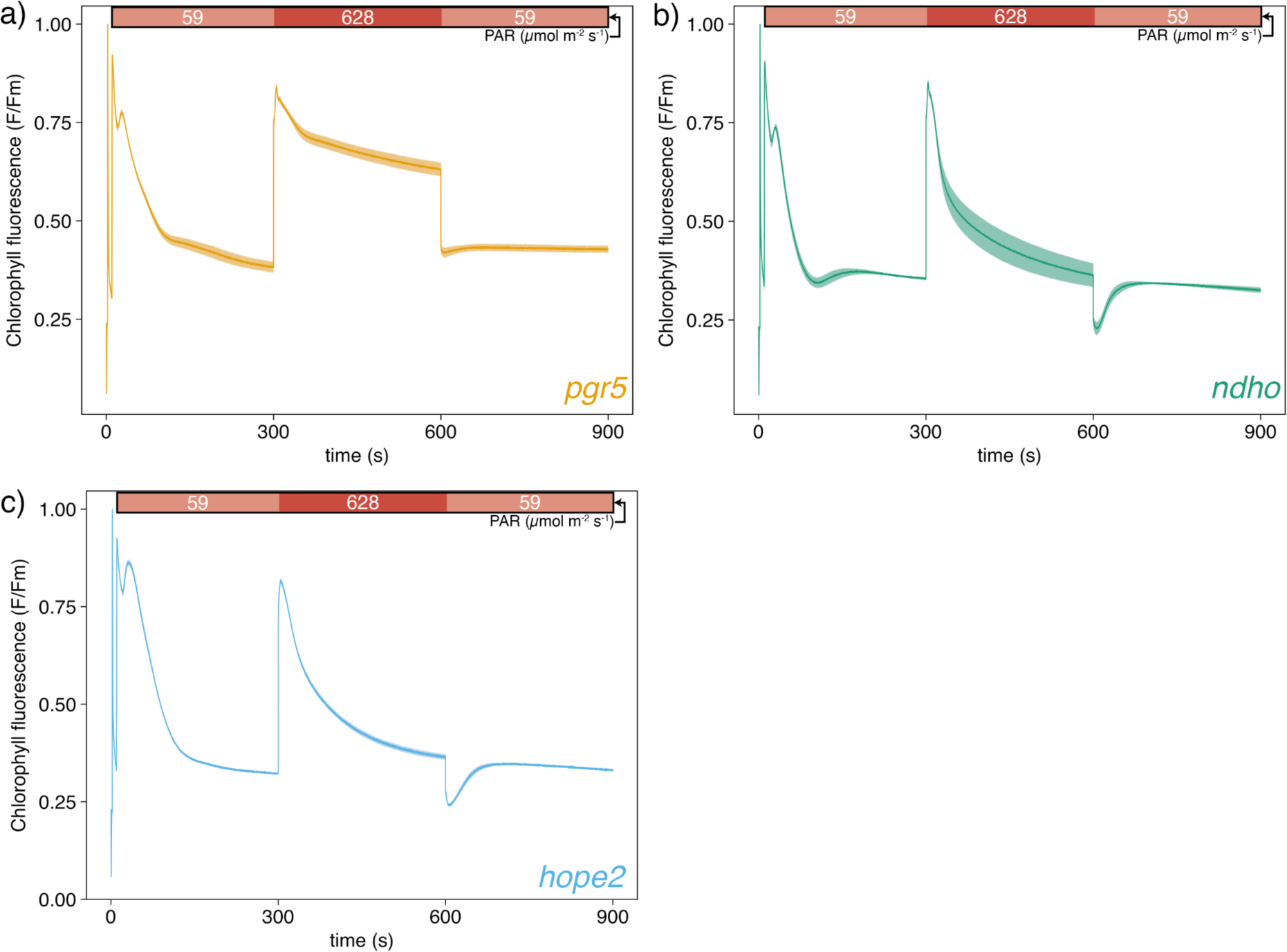
Chlorophyll fluorescence during low-high-low light transition. a) *pgr5*, b) *ndho*, c) *hope2*. Light intensity is indicated at the top of the plot. Fluorescence was normalised to Fm recorded previous to the measurements. Lines show the average of at least three biological replicates and shaded areas represent SEM.

**Supplemental Figure S3:**
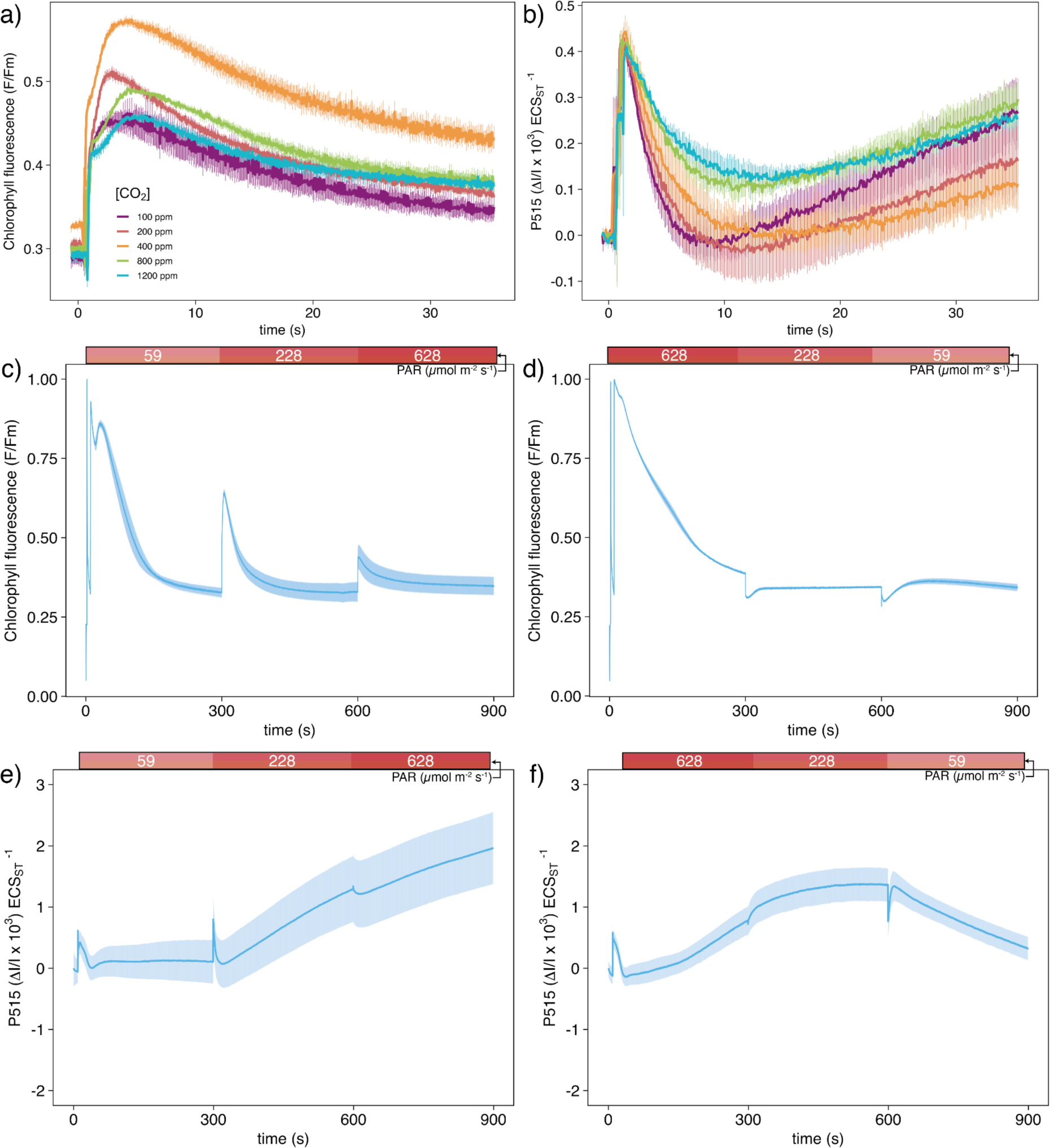
Chlorophyll fluorescence and electrochromic shift signals in *hope2*. a,b) Signals at different CO2 concentrations. c-f) Signals during increasing or decreasing light intensities.

**Supplemental Figure S4:**
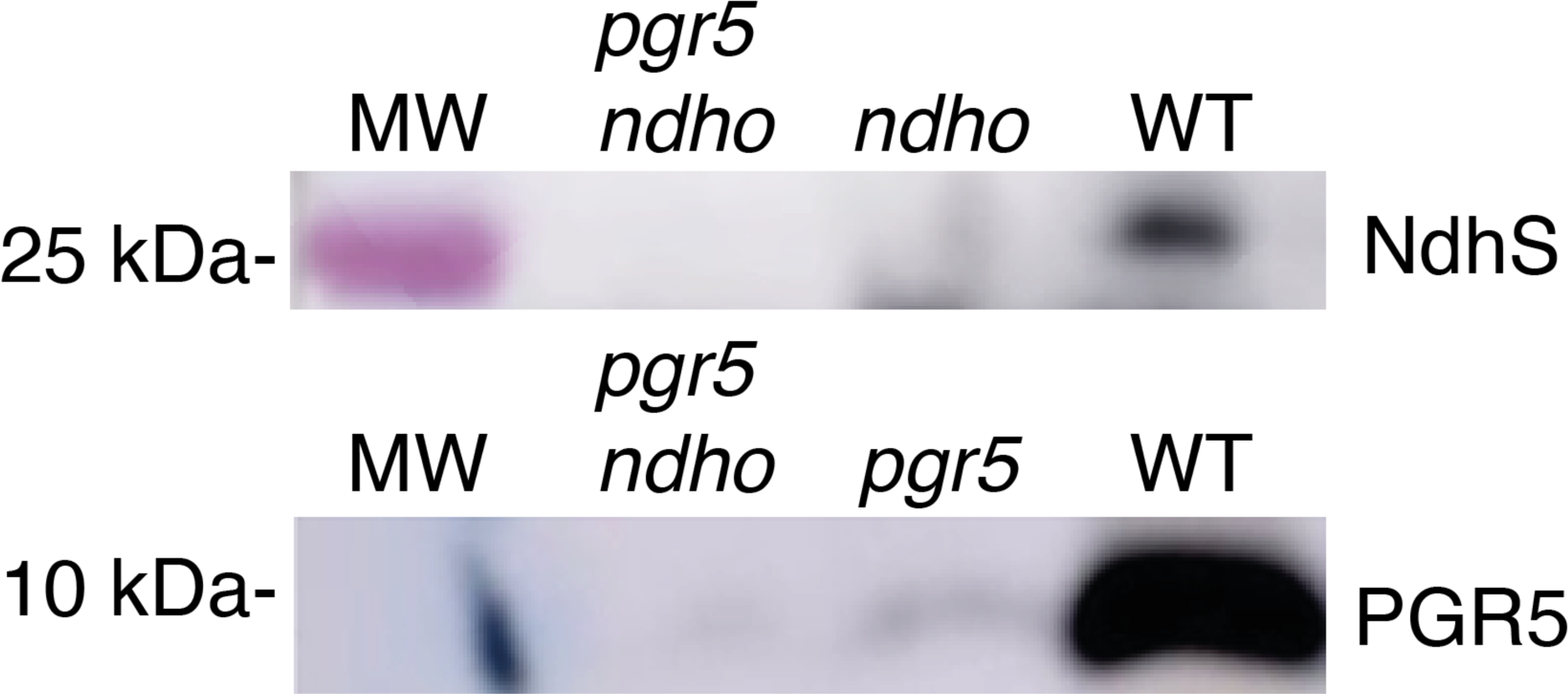
Western blot confirming that *pgr5 ndho* is a complete double knockout of the PGR5 protein and NDH complex.

